# Detecting copy number alterations in RNA-Seq using SuperFreq

**DOI:** 10.1101/2020.05.31.126888

**Authors:** Christoffer Flensburg, Alicia Oshlack, Ian J. Majewski

## Abstract

Calling copy number alterations (CNAs) from RNA-Seq is challenging, because differences in gene expression mean that read depth across genes varies by several orders of magnitude and there is a paucity of informative single nucleotide polymorphisms (SNPs). We previously developed SuperFreq to analyse exome data of tumours by combining variant calling and copy number estimation in an integrated pipeline. Here we have used the SuperFreq framework for the analysis of RNA sequencing (RNA-Seq) data, which allows for the detection of absolute and allele sensitive CNAs. SuperFreq uses an error-propagation framework to combine and maximise the information available in the read depth and B-allele frequencies of SNPs (BAFs) to make CNA calls on RNA-seq data. We used data from The Cancer Genome Atlas (TCGA) to evaluate the CNA called from RNA-Seq with those generated from SNP-arrays. When ploidy estimates were consistent, we found excellent agreement with CNAs called from DNA of over 98% of the genome for acute myeloid leukaemia (TCGA-AML, n=116) and 87% for colorectal cancer (TCGA-CRC, n=377), which has a much higher CNA burden. As expected, the sensitivity of CNA calling from RNA-Seq was dependent on gene density. Nonetheless, using RNA-Seq SuperFreq detected 78% of CNA calls covering 100 or more genes with a precision of 94%. Recall dropped markedly for focal events, but this also depended on the signal intensity. For example, in the CRC cohort SuperFreq identified 100% (7/7) of cases with high-level amplification of ERBB2, where the copy number was typically >20, but identified only 6% (1/17) of cases with moderate amplification of IGF2, typically 4 or 5 copies over a smaller region (median 5 flanking genes for IGF2, compared to 20 for ERBB2). We were able to reproduce the relationship between mutational load and CNA profile in CRC using RNA-Seq alone. SuperFreq offers an integrated platform for identification of CNAs and point mutations from RNA-seq in cancer transcriptomes.

The software is implemented in R and is available through GitHub: https://github.com/ChristofferFlensburg/SuperFreq.

## Introduction

Somatic copy number alterations are important drivers of cancer development[1]. Genomic DNA can reveal genome wide copy number status through different technologies, such as karyotyping[2], SNP-arrays[3], or next generation sequencing of exomes or genomes[4, 5]. Transcriptome sequencing is commonly employed to study gene expression in cancers, but also holds information on the genomic copy number, through its influence on gene expression and variant allele frequencies of heterozygous germline polymorphisms (B-allele frequencies, BAFs). Tens of thousands of public cancer transcriptomes have been sequenced[6], many without matched DNA-sequencing[7], and exploring the somatic copy number alterations may reveal new insights in existing data sets.

Detecting CNAs from RNA-Seq is a challenging task because coverage is highly variable owing to differences in gene expression. Recently, several methods have emerged for calling CNAs from single cell RNA-Seq data[8, 9], and there are to our knowledge currently three publicly available methods to call CNAs from bulk RNA-Seq: CNVkit-RNA[10], CaSpER[11] and CNAPE[12]. However no current approaches accommodate small numbers of samples, paired tumour normal comparisons or absolute copy number calling.

We previously developed SuperFreq to track somatic mutations in cancer, primarily for the analysis of exomes[13]. SuperFreq identifies somatic and germline small variants, calls absolute allele sensitive CNAs and tracks clones across samples. SuperFreq was developed with a methodological focus on empirical variance estimates and error propagation to account for variable sample quality. Here we generalise this algorithm to RNA-Seq with only minor changes in implementation, as the algorithm naturally adapts to the variation in coverage resulting from differences in gene expression.

In this paper, we describe the implementation of the RNA mode for SuperFreq and assess its performance by comparison to array-based copy number calls across two large cancer cohorts. We adopted a strategy that leveraged genes with robust expression across the cohort to improve performance. While SuperFreq represents a significant advance in terms of CNA detection from RNA-Seq, the resolution is lower than DNA array based calls, which has implications for detecting focal events. However, unlike arrays, SuperFreq on RNA-Seq allows for the integrated analysis of CNA calls and point mutations and we reproduce the established relationship between CNA profile and mutation load in CRC. This added functionality opens up the potential to look for more complex types of mutations, including biallelic inactivation of tumour suppressor genes. These results demonstrate how SuperFreq can be leveraged to analyse the mutational landscape of large cancer cohorts from RNA-Seq.

## Methods

The SuperFreq algorithm for identifying SNVs, CNAs and clonal tracking using exomes has been described in detail[13]. The software can now be run in RNA mode, which includes key modifications to the CNA calling algorithm. The algorithm for CNA calling makes use of both read depth and BAFs, but it is mainly the treatment of the read depth that is altered in RNA mode. SuperFreq requires a panel of at least 2 reference normal samples (5-10 recommended) that are used as baseline for read depth. For the analysis of RNA we suggest the reference normal samples are from a highly related tissue and it is ideal that they are processed together with the test samples to limit the batch influence. Exon level read counts are derived for the test samples and the reference normals, then they are corrected for multiple factors such as GC-bias and sex. For each test sample, the corrected pseudo-counts are summed by gene and compared to the reference normals in a one-against-many differential analysis using voom[14] and limma[15]. The output of limma is a log fold change (LFC) *l_i_* of the test sample with respect to the reference normal samples for each gene *i*, as well as a moderated t-statistic *t_i_* that is related to the accuracy of the LFC. SuperFreq defines *W_i_ = l_i_/t_i_*, the scale factor of the limma t-distribution, as a measure of the variance of the LFC in the downstream segmentation of the genome.

A key aspect of the LFC variance estimation in SuperFreq is the neighbour correction. This correction makes use of the observation that neighbouring genes will typically share the same copy number state, that is they should share the same true LFC. Changes in read depth that result from differences in gene expression represent unwanted variation when calling CNAs from RNA-Seq. While we do not know what the shared LFC is at this point of the analysis, we can compare the LFC of each pair of neighbouring genes *i* and *i* + 1, and compare the difference to what is expected from the limma t-distributions with scale factors *W_i_* and *W*_*i*+1_. For this purpose, we define the variance normalised difference *D_i_* between a pair of neighbouring genes *i* and *i* + 1:

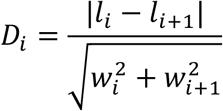

To compare to expected behaviour, we also simulate this difference through

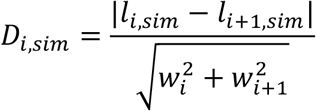

where *l_i,sim_* is a random log fold change drawn from the limma null hypothesis of a moderated t-distribution *l_i,sim_ ~ t*(0,*d*) · *w_i_* where *t*(·) is the t-distribution and *d* is the degree of freedom determined by limma. Note that non-zero but identical LFC of the neighbouring genes cancel out in the numerator of *D_i_*. If the limma variance estimates are accurate, and the assumption that most neighbouring genes share the same LFC holds, then *D_i_* and *D_i,sim_* will be distributed identically, except for a small number of neighbours that span CNA breakpoints. For real data however, the empirical distribution *D_i_* is sometimes wider than *D_i,sim_* for exomes, and almost always wider for RNA-Seq (**Supplementary Figure 1b,d**), indicating that the limma variance estimates *w_i_* are too small. SuperFreq corrects this by adding a constant boost *b* to *w_i_* for all genes *i*:

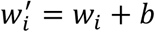

The boost *b* is determined such that the median across genes of the corrected *D′_i_* matches the median of *D′_i,sim_*. Using the median to determine *b* protects from outliers in the empirical distribution, such as neighbours spanning CNA breakpoints. This neighbour correction typically produces a similar shape of *D′_i,sim_* and *D′_i,sim_* for exomes (**Supplementary Figure 1b**), so 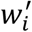 is used as an error estimate of the LFC in the downstream copy number calling. For exomes, the algorithm for LFC error estimation can be summarised as:

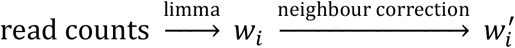

For RNA-Seq data, the tails of *D′_i_* and *D′_i,sim_* are often very different, even with matching medians (**Supplementary Figure 1d**), indicating that the exome algorithm is not adequate for variance estimates of RNA-Seq. To address the incorrect variance estimates, SuperFreq in RNA mode uses the housekeeping gene concept. SuperFreq uses the reference normal samples to identify stable genes; those that are highly expressed with low variance across samples. Genes outside the stable set receive a penalty in the copy number analysis. SuperFreq first selects genes based on mean log read count *q_i_* across the reference normals. Genes with read counts between the 65th and 98th percentile in *q_i_* are selected, corresponding to cut-offs *q_0.65_* and *q_0.98_*. Within these genes, the genes with *w_i_* below the median *w*_0.5_ are selected as stable genes, as shown in **Supplementary Figure 1e.** Genes outside of the stable set are down-weighted based on how much they deviate from the stable range in expression *q* and scale factor *w*: Gene *i* with log read counts *q_i_* outside the range (*q*_0.65_, *q*_0.98_) are down-weighted by adding a penalty Δ_*i,q*_ to *w_i_* of

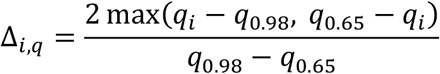

*Δ_i,q_* is limited to a maximum of 1. This penalty down-weights genes depending on how much the mean log read count of the gene in the reference normals differs from the log read counts of the stable genes. Similarly, genes with *w_i_* larger than the cutoff *w*_0.5_ receive a penalty Δ_*i,w*_ to *w_i_* of

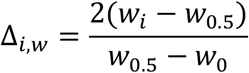

Where *w*_0_ is the smallest *w_i_* across all genes. Δ_*i,q*_ is also limited to a maximum of 1. Δ_*i,q*_ and Δ_*i,w*_ are added to *w_i_*, increasing the variance estimate of all the nonstable genes.

In the following step of the RNA mode algorithm, only the stable genes are used to determine the boost *b* in the neighbor correction, but otherwise the downstream copy number calling algorithm proceeds unchanged. The variance estimate algorithm in RNA mode can be summarized through

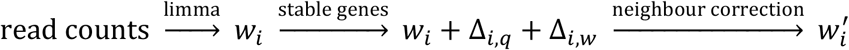

The increased variance estimate in RNA mode effectively downweights the LFC, and the algorithm instead relies more on the information from the BAFs when determining CNAs.

### Assessment strategy

We assessed the performance of the CNA calls from RNA-Seq with SuperFreq by comparing to CNA calls from matched SNP-array data analysed with ASCAT[16]. We analysed the Acute Myeloid Leukemia (AML, 116 samples with matched exome and RNA-Seq)[2] and Colon Adenocarcinoma (CRC, 377 samples)[4] cohorts from The Cancer Genome Atlas (TCGA) to study performance across two different rates of somatic CNAs. We also analysed 415 AML samples from Leucegene[17, 18]. SuperFreq relies upon a set of normal diploid controls; for RNA-Seq it is preferable that these are from the same tissue that gives rise to the cancer. For the CRC cohort, we selected the reference normals from the set of matched normal tissue that was extracted from adjacent colon, which was available for 17 cases. For the AML cohort, matched control tissue was not available, so we used 10 CD34+ sorted cells from the Leucegene cohort as reference normals. These also served as reference normals for the AML samples in Leucegene. The sex chromosomes were excluded from all comparisons, as X-inactivation in females complicates the calculation of BAFs and makes the CNA calls from RNA-Seq unrepresentative of the genomic copy number.

### Computational Performance

SuperFreq typically analysed an 80M reads TCGA CRC RNA-Seq sample against 17 reference normal samples in 2 hours, using 4 cpus and 30GB of memory. Samples can be analysed in parallel with a moderate improvement in per-sample performance due to caching of the reference normal analysis.

## Results

### SuperFreq can detect copy number alterations in RNA-seq

In order to assess the ability of calling copy number alterations in RNA-seq we made use of TCGA samples where both SNP-arrays and RNA-Seq was available. After calling CNAs with SuperFreq on RNA-seq, we compared the results to SNP-array calls from ASCAT. First we compared the coverage LFC and BAF of the RNA-Seq and SNP-array segments. An example CRC sample, TCGA-AD-A5EK, is shown in **Figure 1a**. The SNP-array LFC is calculated relative to a matched normal sample, while the RNA-Seq LFC is calculated relative to a set of RNA samples that act as reference normals. We measure the fraction of the genome of each sample that agreed within 0.15 for LFC and 0.05 for BAF (which represents the average signal from the loss or gain of a single chromosome in 20% of the cells). When comparing CNA calls from RNA-seq to those from ASCAT, the AML samples typically had agreement over more than 80% of the genome (**Figure 1b**, blue violin), with 90% of samples agreeing over 80% of the genome. However, the distribution was bimodal. The LFC estimate is normalised based on ploidy, so a difference in the ploidy estimate produces a disagreement in LFC genome wide. Indeed, agreement between methods increased to 98% when considering samples with consistent ploidy estimates. The CRC samples have a higher rate of CNAs, providing a more complicated LFC and BAF profile to reproduce (**Figure 1b**, red violin). Restricting our analysis to CRC samples with similar ploidy estimates (**Figure 1c** and **Supplementary Figure 2**), we found agreement between SuperFreq and ASCAT across a median of 87% of the genome, with 65% of samples agreeing over 80% of the genome. The high concordance in both AML and CRC shows that RNA-Seq can be used to segment the genome into regions with LFC and BAF that match SNP-array estimates across most of the genome.

**Figure 1:**
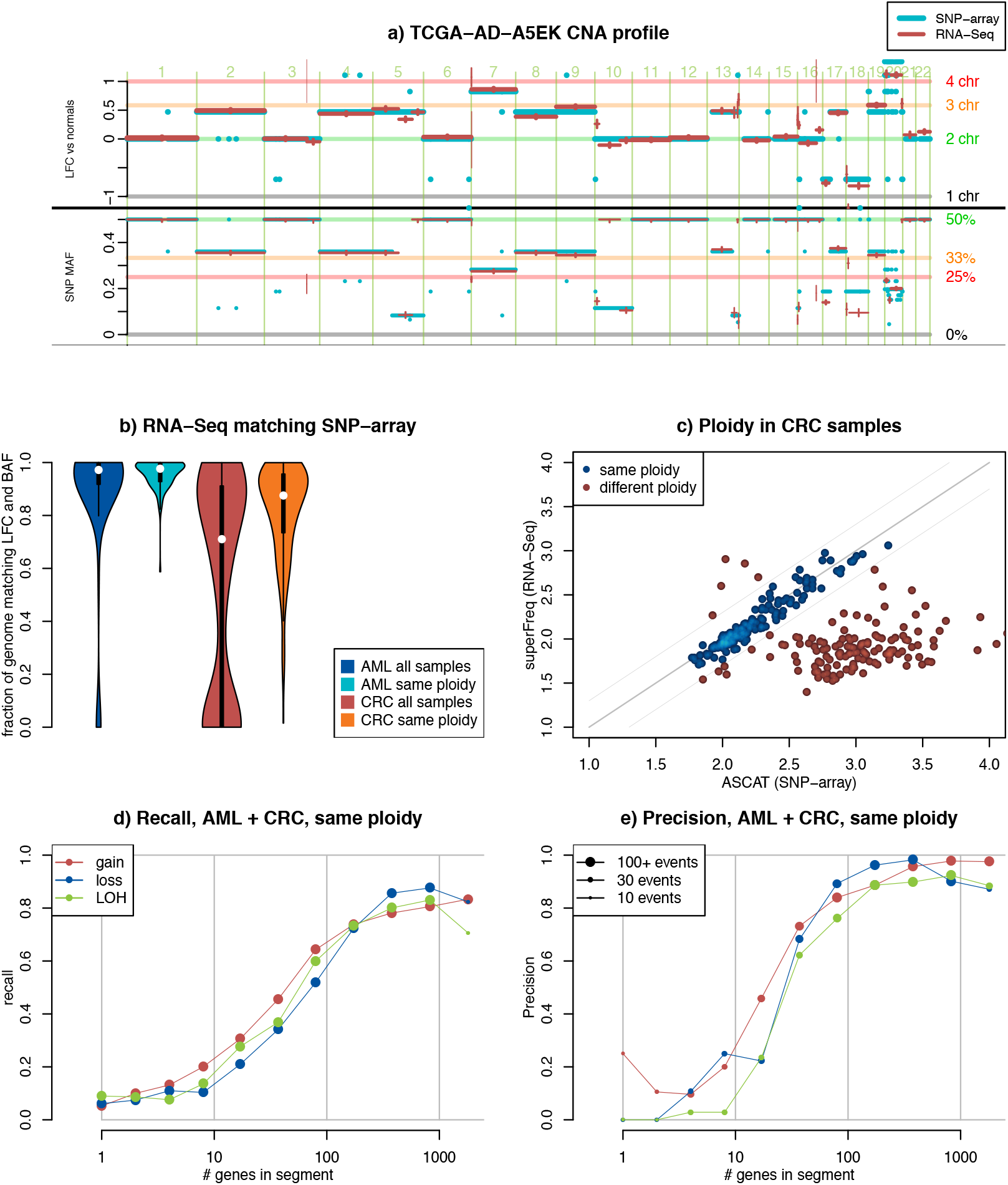
Comparison of CNA calls between SuperFreq on RNA-Seq and ASCAT on SNP-array from TCGA AML (116 donors) and TCGA CRC (377 donors). Sex chromosomes are excluded. a) Copy number call from ASCAT on SNP-array (cyan) and SuperFreq on RNA-Seq (red) from CRC donor TCGA-AD-A5EK. Top panel shows called LFC with respect to the reference normals (SuperFreq) or matched normal (ASCAT), bottom panel shows mirrored BAF. The SuperFreq error bars shows the estimated uncertainty for each segment b) Fraction of the genome for which ASCAT and SuperFreq agrees on LFC within 0.15, and BAF within 0.05. Blue and red violins show AML and CRC samples. Cyan and orange violins show the subset of AML and CRC samples where ploidy agrees within a difference 0.3.. c) The ploidy called by SuperFreq scattered against the ploidy called by ASCAT for CRC. A difference of more than 0.3 as shown by thin grey lines is classified as different ploidy calls as shown by red dots. (AML results are shown in **Supplementary Figure 2**) d) Recall of ASCAT SNP-array calls, split by class of CNA call and number of genes in the segment. Point size indicates the total number of segments. e) Precision of SuperFreq RNA calls compared to ASCAT SNP-array calls, split by class of CNA call and number of genes in the segment. Point size indicate total number of segments.

We assessed the ability to recall regions of altered copy number from RNA-Seq data as a function of the size of the region. To do this, we classified the SNP-array CNA calls into gain (3+ copies), loss (1 or 0 copies) and copy number neutral loss of heterozygosity (CNN LOH, two copies of same allele), and annotated these calls by the number of genes they impacted. We measured the fraction of segments called from SNP-arrays that had a matching segment detected with RNA-Seq with the same classification that covered at least half of the genomic interval. Looking across both TCGA datasets, and restricting the analysis to samples with consistent ploidy estimates, we found 78% overall recall for segments of all three classes covering more than 100 genes (2779/3586 segments) as seen in **Figure 1d**. The results for each cohort are shown separately in **Supplementary Figure 3a-b**. A segment containing 100 genes corresponds to ~12Mbps on average. Sensitivity of CNA calling on RNA-seq dropped to 9% for segments containing fewer than 10 genes (488/5484 segments). For copy number segments of this size, that is less than 1 Mb, even the SNP-array data can be unreliable. Looking at the precision of the SuperFreq calls, we found that 94% (2686/2846) of CNAs identified by SuperFreq over segments covering more than 100 genes were also detected by ASCAT, suggesting a high degree of precision for large events (**Figure 1e**). CNAs identified by SuperFreq that covered less than 10 genes had a precision of only 9% (31/307), indicating that small CNA calls are mostly unreliable. SuperFreq called less than one event smaller than 10 genes per sample on average, giving a low rate of false positives. SuperFreq achieved an overall precision across all CNA sizes of 86% (9537/11028), displaying a robust performance despite the noisy nature of RNA-Seq data. We found that concordance between RNA-Seq and SNP-arrays was generally consistent across genomic regions (**Supplementary Figure 3c**), with higher levels of discordance in telomeres and centromeres.

### Ploidy estimates from SuperFreq

Identifying ploidy from sequencing or SNP-array data is challenging, and orthogonal technologies like cytogenetics or flow cytometric quantification of DNA content provide more reliable estimates. Subclonal CNAs can be mistaken for a genome duplication, and we note that ASCAT assumes a single clone while SuperFreq accommodates subclones. ASCAT calls a ploidy larger than 2.8 for 9 AML samples, while SuperFreq calls a ploidy below 2.1 for all of these samples (**Supplementary Figure 2**). The TCGA-AML cohort has cytogenetic reports for most samples, including 8 of the 9 samples with high ASCAT ploidy estimates. For 7 of these samples the chromosome number is between 37 and 49 chromosomes, consistent with a ploidy close to 2. The remaining sample, TCGA-AB-2813, has two cytogenetic clones, one with 44-45 chromosomes and one with 82-84. This suggests that ASCAT on SNP-arrays has a higher false positive rate for genome duplication that SuperFreq on RNA-Seq. The CRC cohort has a large number of samples where only ASCAT calls a genome duplication, but there are also many samples where both methods identify an elevated ploidy (**Figure 1c**). Without cytogenetic data from this cohort, it is difficult to resolve the true ploidy of the samples.

### Recall of focal amplifications

Many important cancer drivers are activated through amplification, including high level focal amplification[1, 19]. Focal amplification can be difficult to separate from an overexpressed gene in RNA-Seq. To investigate focal amplifications we selected two genes that are recurrently amplified in CRC, the growth factor IGF2 and the cell surface receptor ERBB2[4], and evaluated gene expression and CNA calls from RNA-Seq.

First we identify outliers in expression of IGF2 and ERBB2, as shown in **Figure 2a & 2d**. ASCAT identified focal amplifications (4 copies or more covering at most 15Mbps) in 17 of the 46 samples with elevated expression of IGF2 (**Figure 2b**), while SuperFreq called only one sample with focal amplification using RNA-Seq (**Figure 2c**). For ERBB2 however, the SNP-array data and the RNA-Seq data identified focal amplification in the same 7 cases, out of a total of 8 cases that show high level expression (**Figure 2d-f**). We investigated why there was such a difference in recall between CNA events involving IGF2 and ERBB2. The focal amplifications over IGF2 typically affected a small region (median of 5 genes) and had a median of 4 copies, so recall of around 10% is consistent with our assessment of performance across the TCGA cohorts (**Figure 1d**). The focal amplifications over ERBB2 were significantly larger (median of 20 genes), with a median of 24 copies called. While a gain covering 20 genes has an average recall around 30% in **Figure 1d**, the magnitude of the signal for ERBB2, both in BAF and LFC, allows SuperFreq to recall all 7 focal amplifications. These results demonstrate that sensitivity is not only affected by the number of genes in a segment but also the magnitude of the signal determined by the increase in copy number.

**Figure 2:**
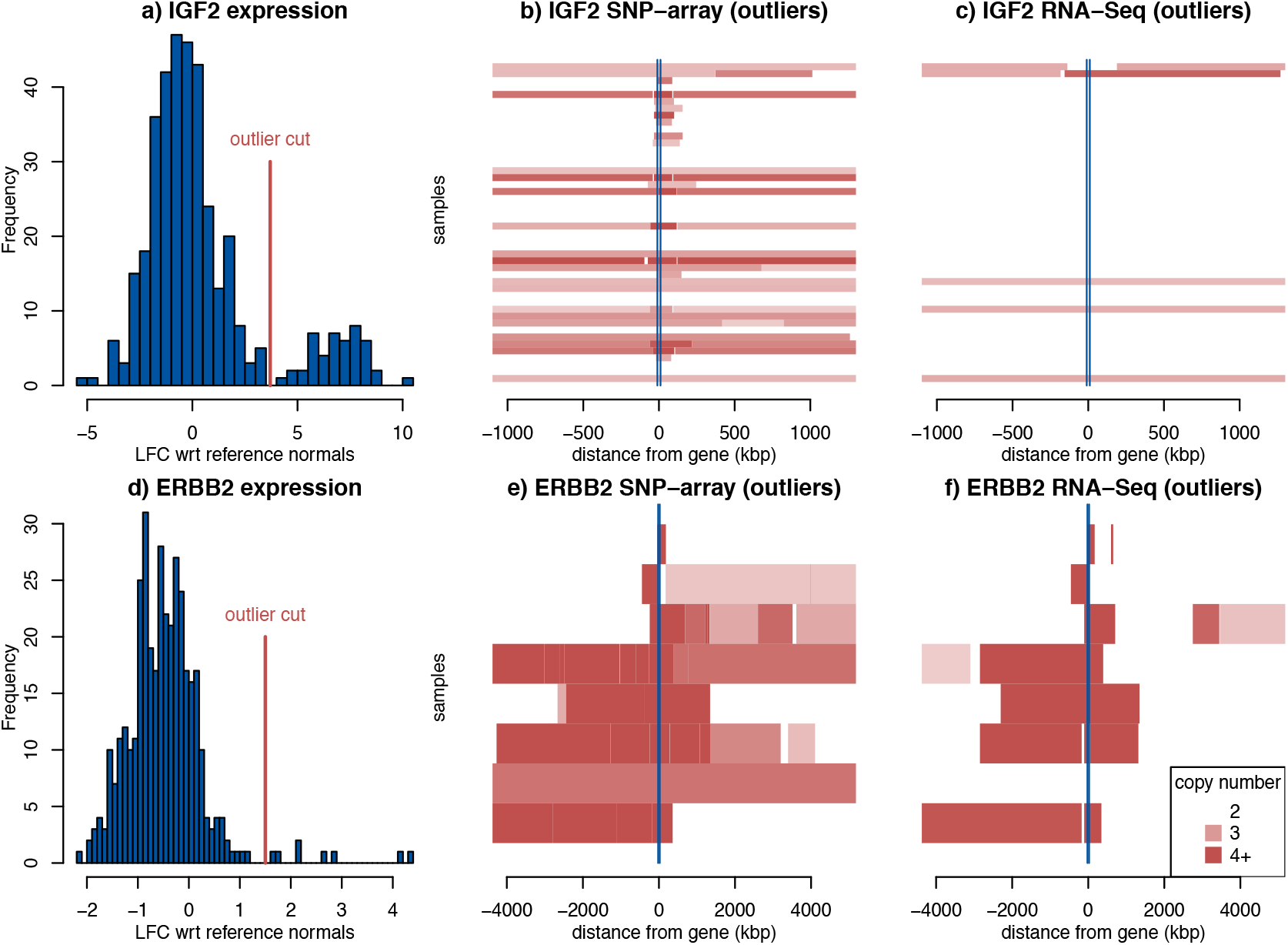
Detection of focal amplifications in TCGA-CRC using RNA-Seq data. a) The expression distribution of IGF2, with a line showing the cut for expression outliers. b) The ASCAT CNA calls around IGF2 for the samples with expression above the outlier cut, ordered with highest expression on top. Opaque red shows increasing amplification. c) SuperFreq CNA calls across the same region of IGF2. The same analysis was performed for ERBB2, and we show d) expression outliers, e) ASCAT CNA calls and f) SuperFreq CNA calls.

### Reproducing cohort-wide results

The first study of TCGA CRC cohort included the discovery that ~16% of cases exhibited hypermutation, owing to genetic changes in the mismatch repair pathway of the proof-reading polymerase POLE[4]. These hypermutated samples tended to have far fewer CNAs. As SuperFreq identifies somatic point mutations, we set out to reproduce this result using only the RNA-Seq data. SuperFreq uses matched normal samples to identify somatic variants, when they are available. For samples that lack a matched normal (360/377 of the CRC samples), the algorithm relies on reference normal samples and population frequencies to filter technical noise and germline variants.

We confirmed a bimodal distribution in number of somatic point mutations detected in RNA-Seq by SuperFreq (**Supplementary Figure 4**), which allowed us to classify hypermutators (>1300 candidate somatic point mutations detected). Sorting the samples by point mutation burden and looking at the CNAs determined by SuperFreq, we see that the hypermutators are depleted of CNAs (**Figure 3a**). We also recovered the pattern of recurrently affected genomic regions in CRC[4], summarised in **Figure 3b**. Many known driving events are recurrent in the cohort, such as loss or CNN LOH of TP53 at chr17p or gain of MYC at chr8q.

**Figure 3:**
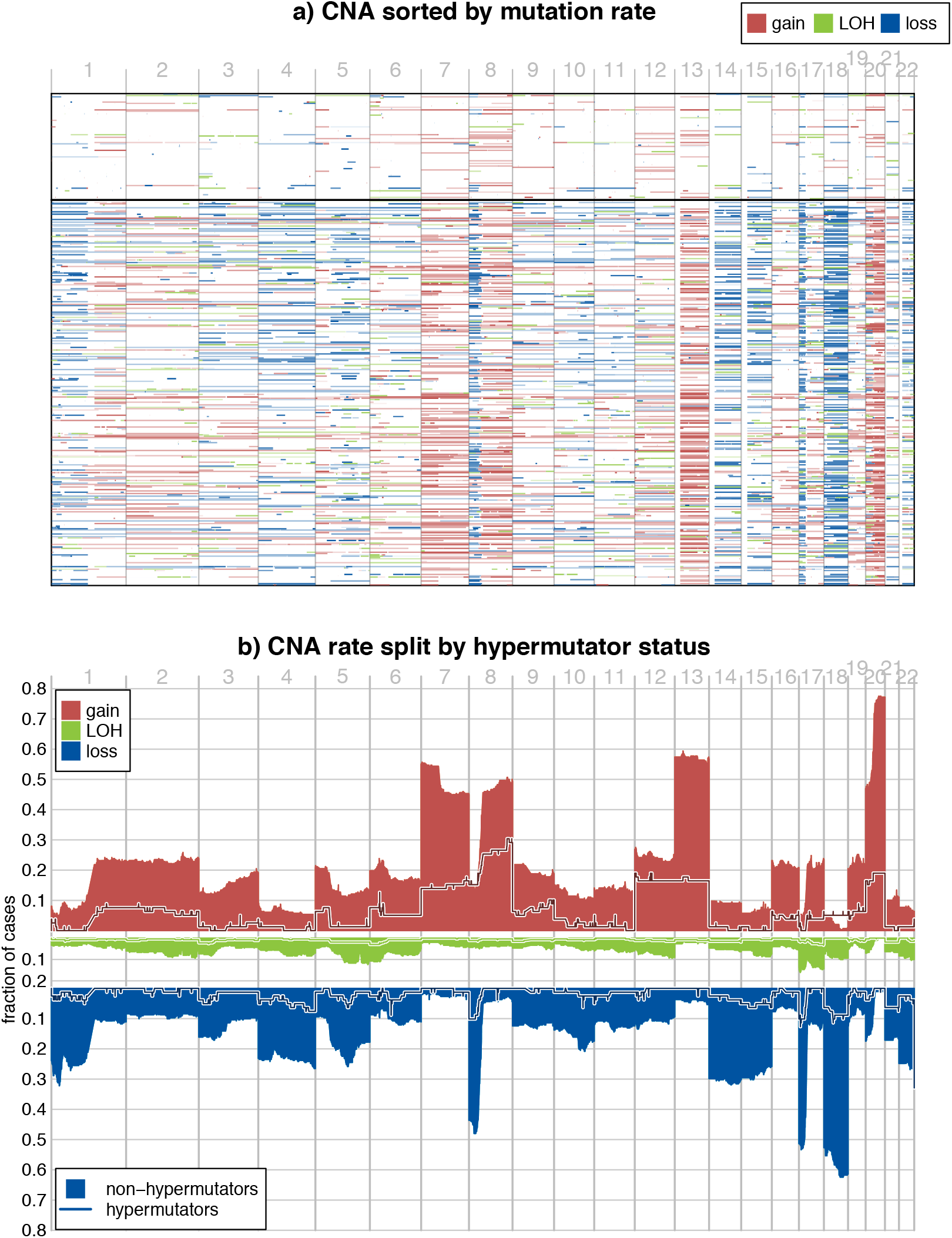
Reproduction of the copy number analysis of the TCGA-CRC cohort using RNA-Seq data. a) The CNA across all TCGA CRC samples sorted by mutation rate with high mutation rate at the top. Red is gain, blue is loss and green is CNN LOH. A black horizontal line marks the hypermutator cut. b) The rate of copy number gain (3+ copies), CNN LOH and loss (one or no copy) across the genome for non-hypermutators (filled shapes), and hypermutators (lines).

We also performed cohort level analysis for the AML samples and provide average CNA profiles for the TCGA and Leucegene cohorts in **Supplementary Figure 5**. While the overall rate of CNAs is significantly lower in AML, SuperFreq still identified established recurrent events, such as monosomy 7, trisomy 8 and loss or CNN LOH of 17p[2].

### Calling point mutations with SuperFreq

Last, we assessed SuperFreq’s ability to call somatic point mutations from RNA-Seq in the CRC cohort. We compared our results to somatic SNVs called from exomes as part of TCGA-CRC, taking variants identified by at least two out of four callers as truth (call sets from mutect2, MuSE, SomaticSniper and Varscan are available through GDC). As mentioned, the mutation rate from RNA-Seq largely reproduces the separation into hypermutated and non-hypermutated samples defined with exomes sequencing (**Supplementary Figure 6a**). APC, TP53 and KRAS are the most mutated genes in CRC, so we evaluated recall of protein coding mutations in these genes. For KRAS and TP53 (median read depth of 29 and 161 respectively), we recalled 79% and 91% of coding variants identified by TCGA, respectively. However, due to low expression of APC (median read depths of 6), we achieved a recall of only 13% of mutations called from the exomes. Across the three genes, we found a recall of 96% when the mean read depth was 30 or larger, but a recall of only 8% when mean read depth was 10 or less (**Supplementary Figure 6b**).

We were interested to leverage the ability to call both SNVs and CNAs to detect biallelic alterations in tumour suppressor genes. To determine whether this was possible, we considered TP53 as a good test case. Using the consensus variants called from exomes, we found 52 of the TCGA-CRC participants had a coding somatic mutation in TP53 with a variant allele frequency (VAF) higher than 70%. The high VAF indicates that the unaltered copy is no longer present in the cancer which can serve as truth for CNA assessment. Indeed, 50 of these 52 participants (96%) had a CNA call over TP53 detected by SuperFreq on RNA-Seq, which consisted of a loss (n = 32), CNN LOH (n = 14) or amplified LOH (n = 4) (**Supplementary Figure 7**). The results with TP53 suggest SuperFreq on RNA-Seq displays good recall, even for CNN LOH, where the signal is only present in the BAFs, and that this approach will be useful for the detection of biallelic alterations in other tumour suppressor genes.

### Discussion

Traditionally RNA-Seq has only been used for quantifying gene expression and detecting fusion genes, but increasingly this data is being leveraged to detect a broader complement of somatic mutations[20]. There are currently very few methods available to call CNAs from RNA-Seq. The paucity of methods reflect the challenges associated with detecting CNAs with highly variable coverage and the smaller number of heterozygous SNPs available for genotyping. In this paper we demonstrate that SuperFreq can be used to detect allele sensitive CNAs from RNA-Seq. When compared to other recently published methods, such as CNVkit-RNA[10], SuperFreq represents a major advance as it provides absolute, allele sensitive and subclone aware CNA calls.

We benchmarked SuperFreq on the AML and CRC cohorts from TCGA by comparing the results to matched SNP-arrays. When ploidy estimates were consistent, SuperFreq recalled 78% of alterations covering >100 genes detected with SNP-arrays, but had low sensitivity when the CNA impacted <10 genes. SuperFreq had a low false positive rate, with 86% precision across CNA calls of all size, and 94% for CNAs covering >100 genes. This demonstrates that while using RNA-seq to call CNAs can miss some events in gene poor regions, the events that are called are highly accurate. Accurately determining the ploidy of a sample is a difficult problem for both RNA-Seq and SNP-array data and needs to be studied further with orthogonal data. SuperFreq ploidy calls from RNA-Seq were largely consistent with those from ASCAT for the AML cohort, where 91% of estimates differed by <0.3, but the level of agreement dropped to 61% when considering the CRC cohort. This difference may be because SuperFreq is more conservative than ASCAT for calling high ploidies, which we also saw when using SuperFreq to analyse exomes[13]. SuperFreq accurately reproduced the low rate of genome duplication in karyotyping data from TCGA AML, which suggests it has a lower false positive rate than ASCAT, but there is a lack of adequate truth data sets to accurately assess performance in cancers with high rates of aneuploidy.

Chromosome wide CNAs can impact the expression of thousands of genes and are easily detected, but as we focus in on smaller regions the signal becomes more variable. This variability can be due to technical differences, but can also result from differences in gene expression between the test sample and the reference normals. To counter this variability we implemented a housekeeping gene framework that assigns reduced weight to highly variable genes, and we used weighted averages across segments to detect changes in LFC. These approaches reduce the number of false positives, but can also limit the sensitivity to focal CNAs. To assess how the algorithm worked for focal amplifications in practice, we examined ERBB2 and IGF2, two genes that are recurrently amplified in CRC. SuperFreq reliably detected amplification of ERBB2, but showed poor performance in detecting amplification of IGF2. This difference in performance likely reflects the different characteristics of the amplified regions; the ERBB2 region tended to be highly amplified (median 24 copies) and covered a moderate number of genes (median 20 genes), whereas IGF2 amplicons had lower copy numbers (median 4 copies) and impacted fewer genes (median 5 genes). A targeted analysis of expression changes for established cancer genes may offer improved sensitivity for key focal amplifications, but extending this approach genome-wide is problematic, as it would likely increase the number of false positive calls.

The choice of reference normals is crucial for accurate CNA calls from RNA-Seq with SuperFreq. In particular it is important that the cell type matches the test samples as closely as possible. While batch effects between different data sets are a potential issue, we successfully used reference normals from Leucegene to analyse the TCGA-AML cohort, showing that reference normals from independent public data sets can be used if required. As well as variable regions like the HLA and immunoglobulin loci, CNA calling from RNA-Seq is also unreliable on the sex chromosomes due to X-inactivation. RNA-Seq of cancer samples are typically performed without a matched normal, which impacts on the quality of both CNA and somatic SNV calling. For SNV calling, absence of a matched normal typically results in an increased false positive rate, due to an inability to remove rare germline variants. Whereas for CNA detection, the absence of matched normal tissue will decrease the sensitivity to detect LOH in cancer samples with high purity. Leveraging variant call sets from matched DNA samples may be one way to improve detection of LOH, or strategies could be implemented to detect regions in which heterozygous germline variants are depleted.

SuperFreq was developed as an integrated analysis pipeline for exomes, and includes identification of somatic SNVs, which allows for an integrated analysis of point mutations and CNAs. We used this approach on RNA-seq to verify the finding that hypermutated CRC cases carry relatively few CNAs. When considering individual variants, SuperFreq displayed good recall of point mutations in expressed genes (95% recall when read depth > 30), but genes with low expression provided little sensitivity for point mutations. These differences in expression level have important implications for detecting mutations in driver genes. Even considering this limitation, combined detection of point mutations and CNAs can help to reveal biallelic loss of tumour suppressors. As an example, we looked in detail at cases in TCGA CRC with somatic mutations in TP53 with a high VAF in the DNA, and found that 50/52 (96%) had a loss or LOH called by SuperFreq in the RNA-Seq data. This suggests SuperFreq can be implemented to detect tumour suppressor genes in large cohorts using RNA-Seq, as long as the gene is expressed at sufficient levels.

SuperFreq provides a robust framework for CNA calling from RNA-Seq. The analysis can be adapted to include a matched normal sample, or to rely on a pool of reference normal samples. The generation of absolute allele aware CNA calls represents a major advance over existing approaches. We provide proof-of-concept that existing RNA-Seq datasets that have only been used for expression analysis can be used for broader mutational profiling to provide new insights into the key molecular events that drive these cancers.

## Availability

SuperFreq is available as an R package on github: https://github.com/ChristofferFlensburg/SuperFreq/ Data and code to reproduce the figures are available at: https://gitlab.wehi.edu.au/flensburg.c/SuperFreqRNApaper

## Acknowledgements

This work was supported by the Australian National Health and Medical Research Council (NHMRC) (Project Grant to IJM 1145912; Independent Research Institutes Infrastructure Support Scheme grant 9000220), the Cancer Council Victoria (grant-in-aid to IJM 1124178), a Victorian State Government Operational Infrastructure Support (OIS) grant; a Victorian Cancer Agency fellowship (to IJM) and the Felton Bequest. We wish to acknowledge the generous support of Mr. Malcolm Broomhead who provided philanthropic support. The funders had no influence over the final content of the manuscript.

The results here are based upon data generated by the TCGA Research Network: http://cancergenome.nih.gov/. GDC provides an invaluable resource for cancer genomics research. Data from TCGA was accessed through a data request through dbGAP and usage is in accordance with our project aims. We also wish to acknowledge the Leucegene consortium for access to sequencing data from primary AML samples (http://leucegene.ca).

## Supplementary materials

**Supplementary Figure 1:**
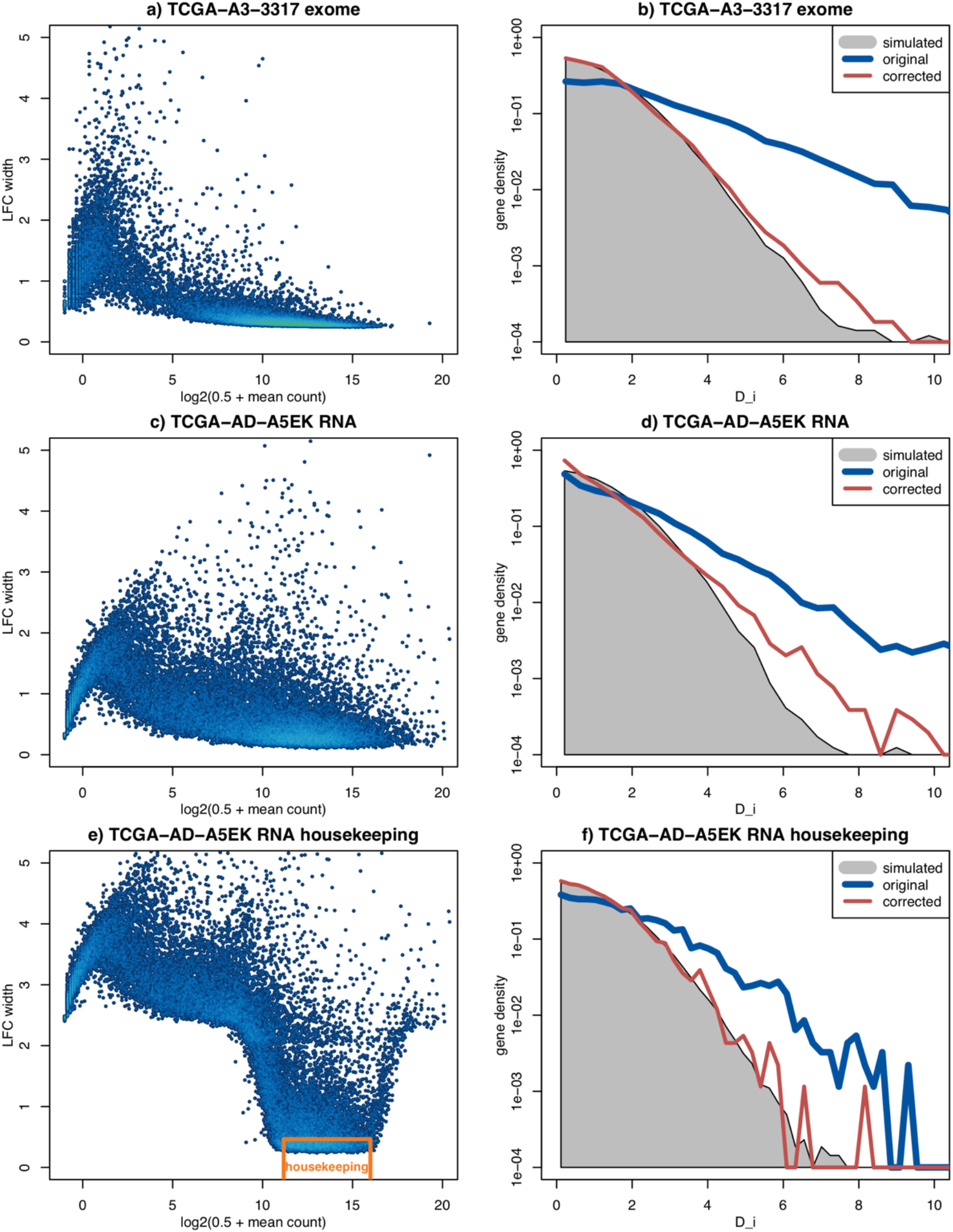
Housekeeping and neighbour corrections. Left plot shows the width of the t-distribution used to approximate the error in LFC compared to the reference normals. Right panel show the desired simulated distribution of Dij (grey), and the distribution of the data before (blue) and after (red) neighbour correction. First row is a low quality exome from TCGA-A3-3317. Second and third row are the RNA-Seq sample from TCGA-AD-A5EK, with third row including housekeeping penalty for the variance estimates, and neighbour correction performed on the stable genes only.

**Supplementary Figure 2:**
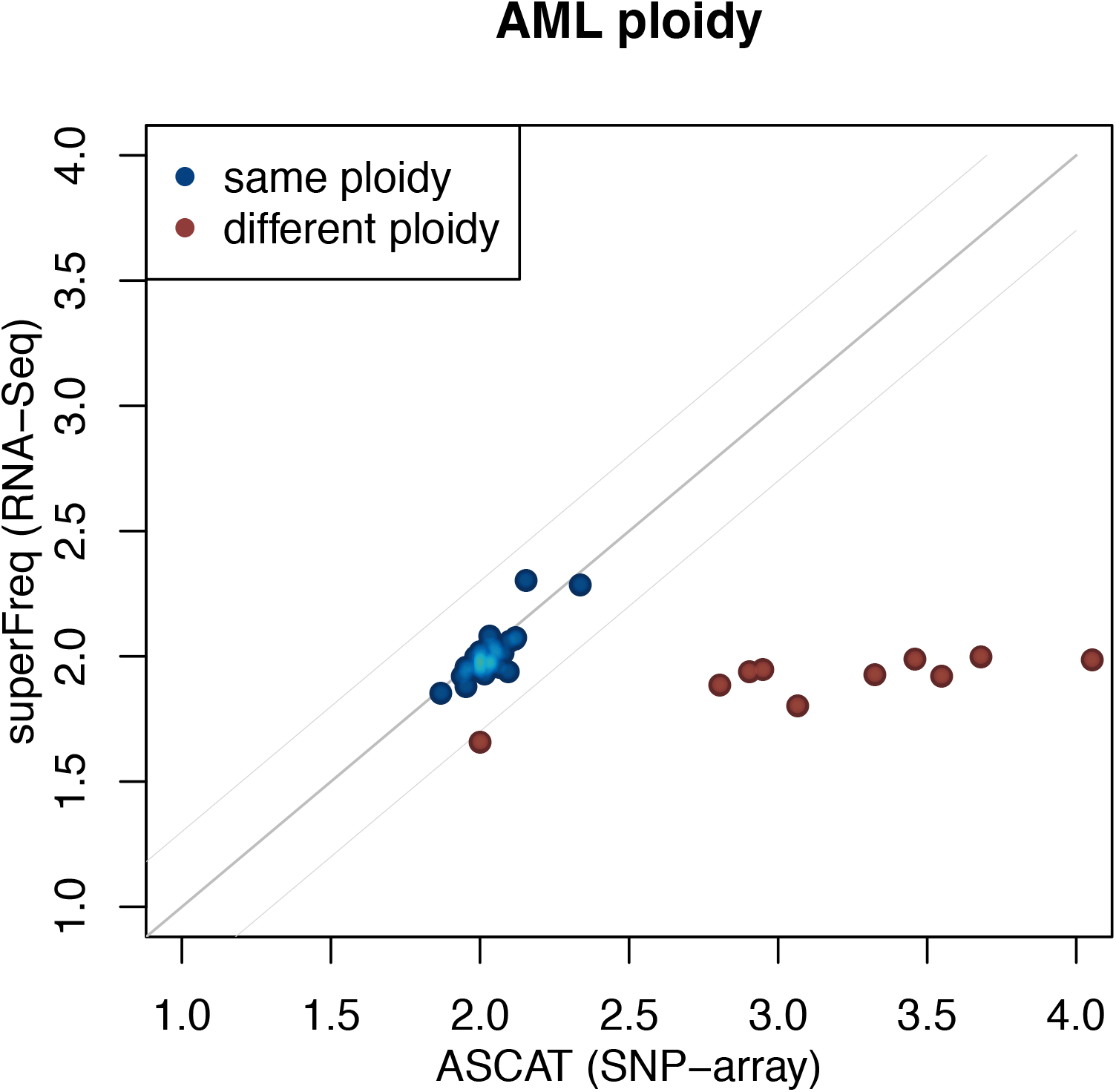
The ploidy called by SuperFreq scattered against the ploidy called by ASCAT for TCGA AML. A difference of more than 0.3 as shown by thin grey lines is classified as different ploidy calls as shown by red dots.

**Supplementary Figure 3:**
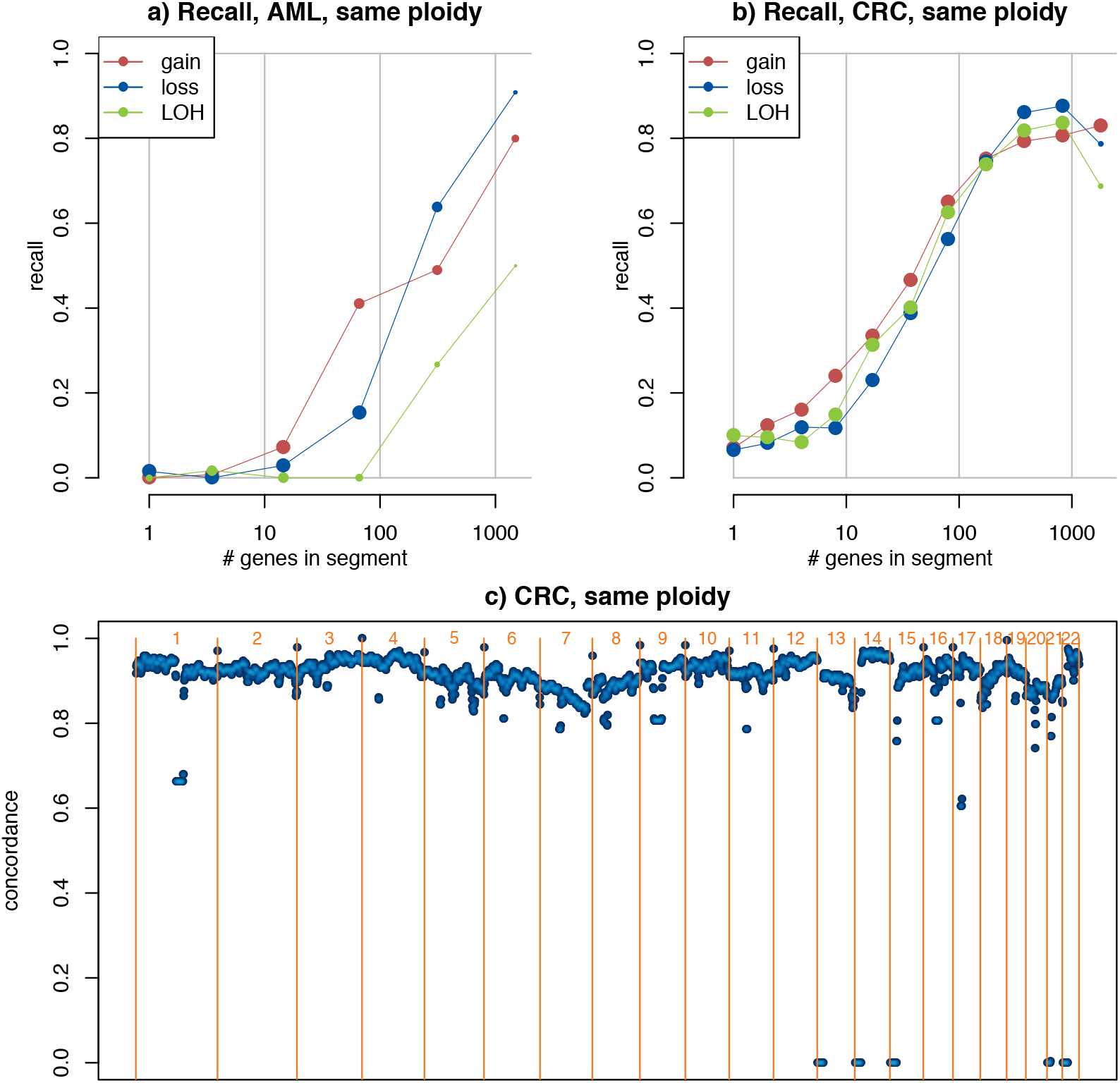
a-b) Recall of copy number alterations for the AML and CRC cohorts. The bin size of the AML data is increased, due to lower sample number and the lower rate of CNAs. c) Classification concordance across the genome for TCGA CRC samples agreeing on ploidy. The samples are classified into gain, loss, CNN LOH and normal as in **Figure 1b**. Y-axis shows the fraction of samples where ASCAT and SuperFreq agrees on classification at the genomic coordinate on the x-axis.

**Supplementary Figure 4:**
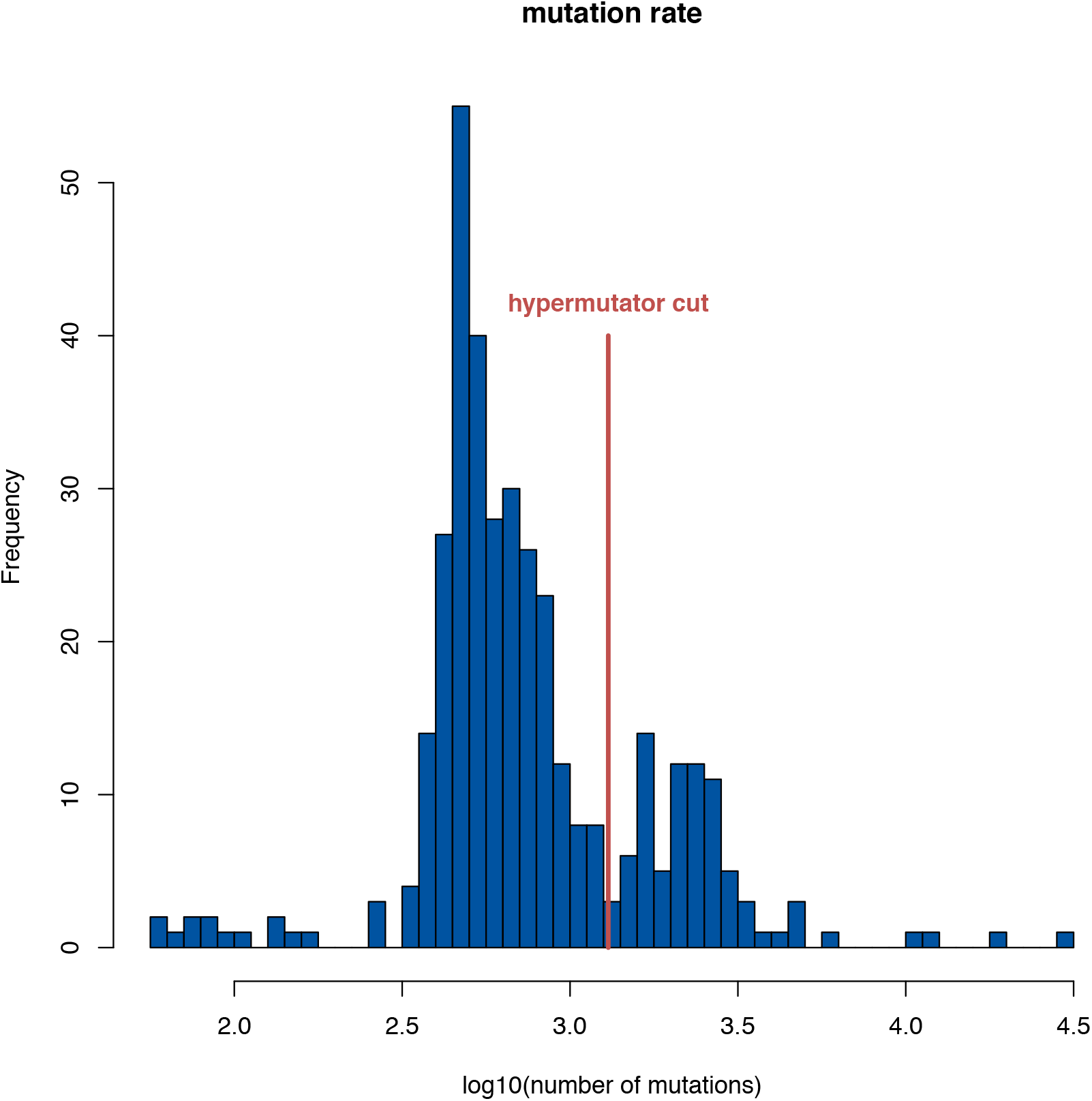
The distribution of mutation rate across the TCGA CRC cohort, as called from RNA-Seq with SuperFreq. Also shown is the cut for hypermutators used in the CNA plots in Figure 2.

**Supplementary Figure 5:**
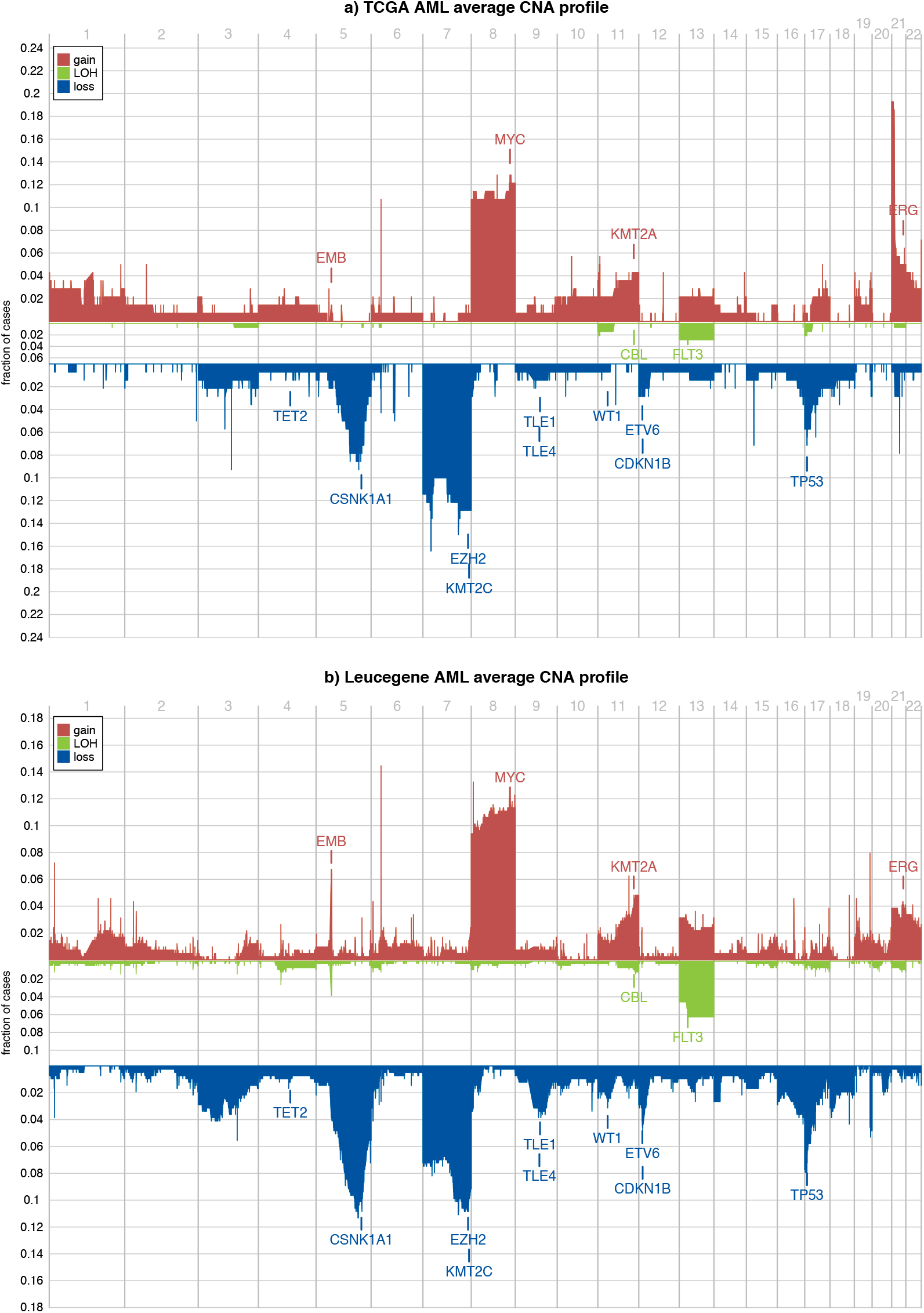
The rate of copy number gain (3+ copies), CNN LOH and loss (one or no copy) in a) TCGA-AML (116 samples) and b) the Leucegene AML cohort (415 samples). Key genes potentially driving selection of recurring CNAs are highlighted.

**Supplementary Figure 6:**
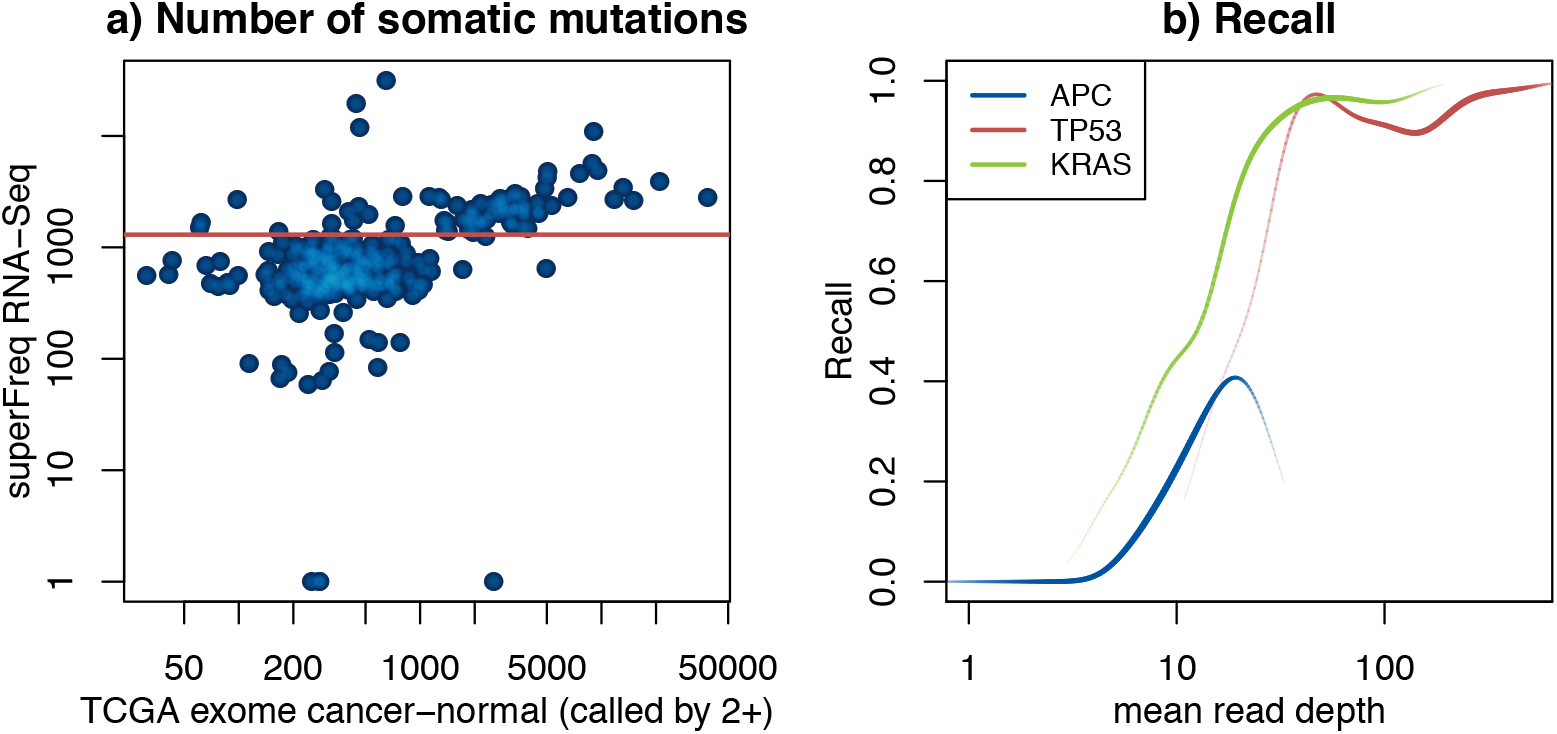
a) The number of somatic point mutations called on TCGA CRC by SuperFreq from RNA-Seq against the number of point mutations called by at least two of the four methods by TCGA on WXS. The cut for hypermutator in the SuperFreq mutation rate is shown in red. The spearman correlation is 0.46. b) Recall of coding mutations in RNA-Seq as a function of mean read depth across the gene. Variants called by at least two methods in matched exomes are used as truth. Recall has been smoothed and shown for the three most mutated genes in the TCGA CRC cohort, where line thickness indicates number of variants available as truth. Mean read depth of 10 corresponds to 1 FPKM in a 50M 2×100 paired end library.

**Supplementary Figure 7:**
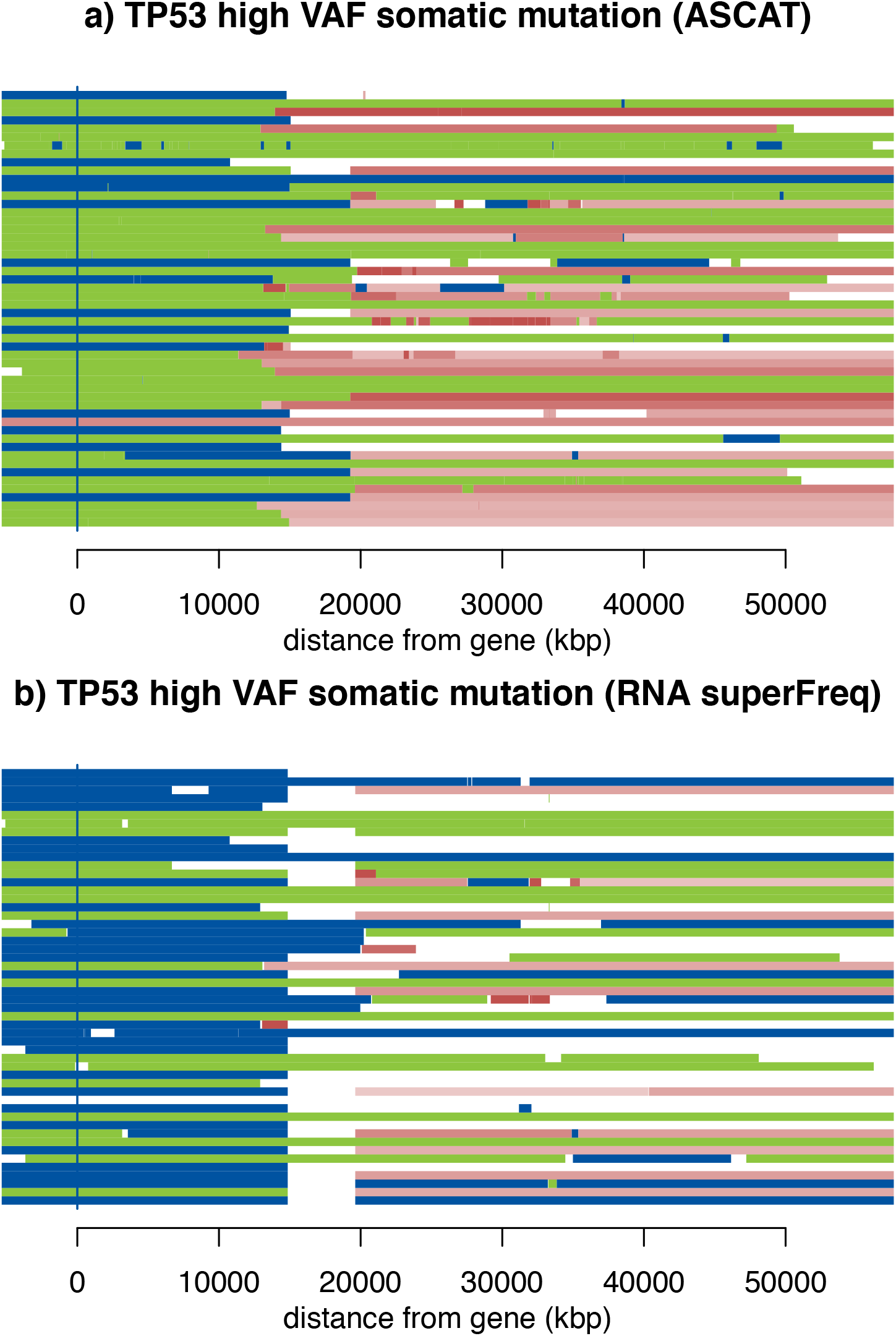
Copy number calls across chromosome 17 for 52 TCGA CRC patients with a somatic TP53 mutation with VAF > 70% called from matched exomes. Blue indicates loss, green indicates LOH with 2 or more copies, and red indicates gain without LOH. Copy number call from ASCAT on SNP-array (a) and SuperFreq on RNA-Seq (b). Due to different ploidy calls, many samples have a loss called by SuperFreq, but CNN LOH called by ASCAT, both are plausible mechanisms for biallelic inactivation and elevated VAF.

